# Patterns of tumor progression predict small and tissue-specific tumor-originating niches

**DOI:** 10.1101/330175

**Authors:** Thomas Buder, Andreas Deutsch, Barbara Klink, Anja Voss-Böhme

## Abstract

Cancer development is a multistep process in which cells increase in malignancy through progressive alterations. The early phase of this process is hardly observable which aggravates an understanding of later tumor development. We shed light on this initial phase with a cell-based stochastic model calibrated with epidemiological data from the tissue scale. Our model allows to estimate the number of tumor cells needed for tumor formation in human tissues based on data on the diagnosed ratios of benign and malignant tumors. We find that the minimal number of cells needed for tumor formation is surprisingly small and largely depends on the tissue type. Our results point towards the existence of tumor-originating niches in which the fate of tumor development is early decided. Our estimate for the human colon agrees well with the size of the stem cell niche in colonic crypts. Our estimates might help to identify the tumor-originating cell type, e.g. our analysis suggests for glioblastoma that the tumors originate from a cell type competing in a range of 300 - 1900 cells.

**Summary:** We estimate the number of tumor cells needed for tumor formation in human tissues and propose the existence of small and tissue-specific tumor-originating niches which might help to find tumor-originating cell types, in particular in glioblastoma.

## Introduction

Cancer development is a multistep process in which cells acquire a certain number of progressive epigenetic and genetic alterations [1]. This multistep process can be divided into a neutral and a selection phase. In the neutral phase, the epigenetic and genetic alterations do not confer a proliferative fitness advantage to the tumor precursor cells whereas cells gain such an advantage in the selection phase [2,3]. A single genetically altered cell does not necessarily induce tumor formation but is rather exposed to competition with its corresponding wild-type cells [4]. The realization of this competition depends on the tissue and cell type. It can be direct due to replacement of a cell by the offspring of another proliferating cell [4] but also indirect, for example by symmetric and asymmetric division of cells [5]. Importantly, tissues are composed of different types of cells but only those which are capable to give rise to a progeny able to accumulate alterations can be tumor-originating cell types [6]. Tumor-originating cell refers to the wild-type cell of a certain type that acquires the first alteration in the multistep process of cancer development. Within the neutral phase, the progeny of the tumor-originating cell competes with wild-type cells within normal tissue homeostasis. Because this competition is controlled by the original tissue organization, the range of this competition is determined by the tissue structure which provides natural spatial boundaries for the spread of the progeny of the tumor-originating cell [7,8].

In order to induce tumor formation, the progeny of the tumor-originating cell must not go extinct but has to establish within the tissue. This establishment is achieved by clonal expansion to a sufficiently large cell population [9]. For some tissues, there is experimental evidence that this establishment is characterized by an outcompetition of wild-type cells within the homeostatic range of competition. For example, the tumor-originating cell within the human colon has been identified to be almost always a stem cell with a first hit in the APC gene, and a second hit in this gene is sufficient to induce adenoma formation, a benign precursor of malignant adenocarcinoma. These stem cells reside at the bottom of so-called niches within colonic crypts and are capable of self-renewal and multilineage differentiation [10]. It has been demonstrated that tumor-originating cells neutrally compete with wild-type stem cells for a position within the spatially restricted stem cell niche [5]. Either such an altered stem cell goes extinct due to this competition or eventually replaces all wild-type stem cells within the stem cell niche. This process has been termed monoclonal conversion and represents almost always the first step of tumor formation within the human colon [10]. Hence, the monoclonal conversion of the stem cell niche by the progeny of the tumor-originating cell with loss of the APC gene induces the establishment of an adenoma on the tissue scale. However, in other tissues the early phase of tumor development on the cellular scale is less understood. The main reason is a lack of knowledge regarding the tumor-originating cell type. Similar to the colon, it has been shown that stem cells within the hematopoietic system represent the tumor-originating cell type [11,12]. In contrast, there is also evidence that non-stem cells can be the tumor-originating cell type, e.g. in oligodendroglioma [13]. Although the lineage in which cancer originates has been revealed for skin, pancreatic, brain and breast tumors, the tumor-originating cell type remains elusive in most cases [14]. Its identification may allow earlier detection of malignancies and may lead to preventive therapies for individuals at high risk of developing cancer [14].

On the tissue scale, one observes different types of tumor progression. Tumors can progress sequentially, i.e. with a clinically detectable benign precursor stage. Alternatively, they can also progress by tunneling without such a prior benign precursor stage. Epidemiological data allow to infer the progression patterns with respect to the ratios of tunneling versus sequential progression of different tumors. Interestingly, these progression patterns differ largely between tissues. Some tumors exhibit predominantly sequential progression, e.g. benign adenoma almost always develop prior to adenocarcinoma in the colon [10]. Similarly, multiple myeloma are in almost all cases preceded by a premalignant state called monoclonal gammopathy of undetermined significance (MGUS) [15]. In contrast, glioblastoma develops in 90% of all cases without evidence of a less malignant precursor lesion (primary glioblastoma) and progresses in 10% of all cases from low-grade tumors (secondary glioblastoma) [16]. In which way these progression patterns on the tissue scale emerge from the multistep process of cancer development on the cellular scale is difficult to infer since the early phase of this multistep process is hardly observable.

In this work, we use observables on the tissue scale to shed light on the early cellular processes of tumor development. We utilize a Moran model with mutations [17–19] to describe cellular competition between wild-type cells and tumor cells. Benign and malignant tumor subtypes on the tissue scale are represented by two absorbing states within the model. We incorporate epidemiological data on the progression patterns of cancers to calibrate the model. By analyzing the model dynamics with respect to different spatial cell arrangements, we obtain a lower and upper bound for the critical number of tumor cells needed for tumor development on the tissue scale. Interestingly, our estimates are considerably small, tissue-specific and far away from the overall number of cells in a clinically observable tumor. We therefore propose that the fate of tumor development is decided in tissue-specific tumor-originating niches. This proposal is supported by our estimate of the tumor-originating niche size for the human colon which agrees well with the size of the stem cell niche in colonic crypts. In particular, we propose that a tumor-originating niche size of 300 – 1900 cells within the human brain can explain the ratio of primary and secondary glioblastoma. Interestingly, our estimates also agree well with the minimal number of tumor cells needed for tumor formation in mice injection experiments and might allow to infer the tumor-originating cell type.

## Materials and methods

### State space and representation of benign and malignant tumor subtypes

The multistep process in which cancer cells increase gradually in malignancy differs with respect to the number of steps, e.g. two steps in retinoblastoma [20] compared to seven steps in colon cancer [21]. In our cell-based model, we only regard the last step within the neutral phase and the first step within the selection phase such that we obtain a two-step process. This coarse-grained approach is appropriate for our purpose since we are only interested in modeling tumor progression patterns and not quantities which are largely influenced by the precise number of steps, e.g the time-scale of tumor development or intra-tumor heterogeneity. In the cellular two-step process, genetic or epigenetic alterations can transform *wild-type cells* into *benign tumor cells* which can further progress to *malignant tumor cells*. We assume that the benign progeny of the tumor-originating cell competes with wild-type cells and can clonally expand within normal tissue homeostasis. The parameter *N* in our model describes the homeostatic range of this competition. We further assume that a benign tumor on the tissue scale will develop if monoclonal conversion of wild-type cells by benign tumor cells within the homeostatic range of competition *N* is achieved. This assumption is based on experimental observations for example within the colon where mutant cells either go extinct or fixate in the colonic stem cell niche [5]. Moreover, mice injection experiments indicate that a critical number of tumor cells is needed for tumor formation [22–29] which suggests a point of no return on the cellular scale for tumor formation. If the first benign tumor cell progresses to a malignant tumor cell we assume that a malignant tumor will inevitably develop on the tissue scale which reflects a high fitness advantage of malignant tumor cells.

The state space of the underlying stochastic process of the model is *S* ={ 0, 1, 2, *N, E*}where states 0 to *N* represent the occurrence of the respective number of benign tumor cells without the occurrence of malignant tumor cells. State *E* indicates the presence of a malignant tumor cell. States *N* and *E* correspond to emergence of benign and malignant tumor subtypes and therefore to sequential and tunneling tumor progression, see also Figure1. Both states *N* and *E* are absorbing states of the underlying stochastic process, see also Text S1 for details.

**Figure 1:**
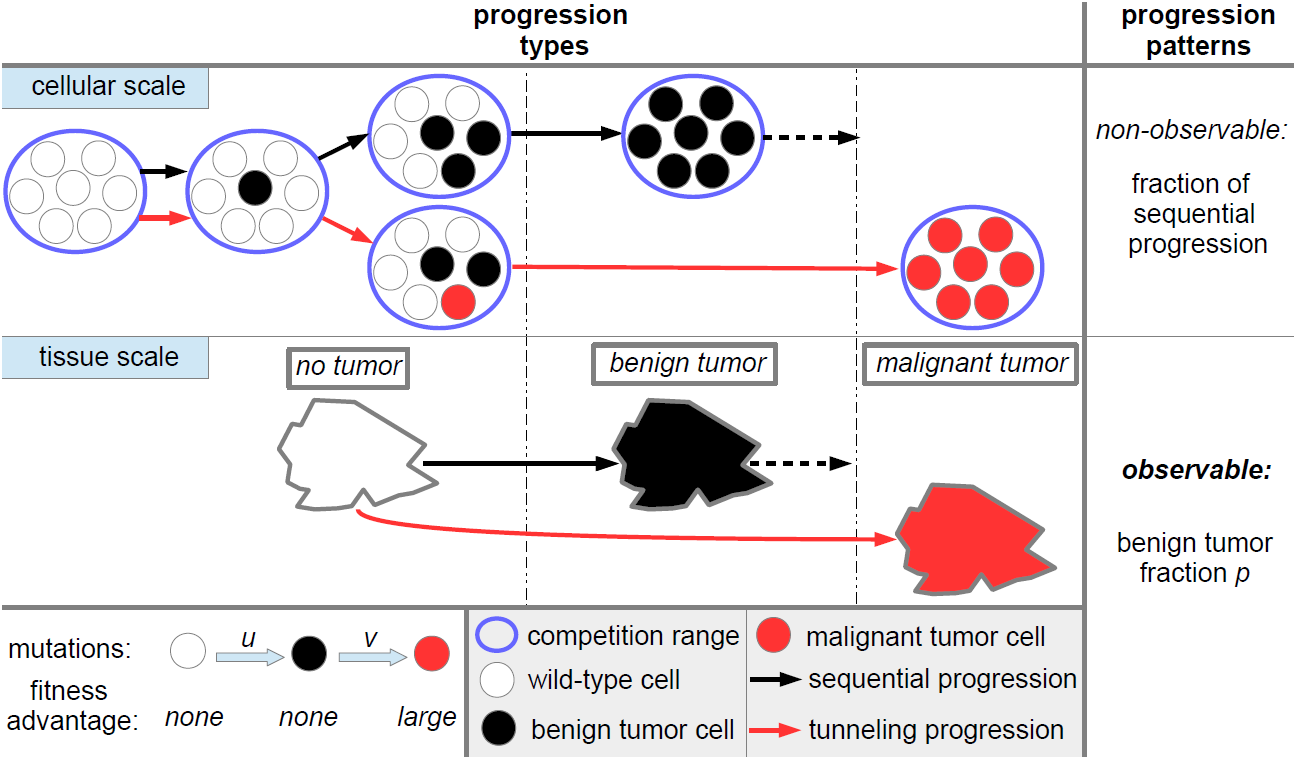
Tumor progression types and patterns in the model. On the cellular scale, wild-type cells can progress to benign tumor cells during proliferation with mutation probability *u*. Wild-type cells and benign tumor cells neutrally compete with each other within the homeostatic range of competition which is modeled by Moran dynamics, see Figure 2. A benign tumor will manifest if all cells within the homeostatic range of competition are converted to benign tumor cells. Furthermore, benign tumor cells can progress to malignant tumor cells during proliferation with probability *v*. Then, a malignant tumor inevitably develops. These cellular dynamics lead to two distinct progression types at the tissue scale, namely sequential progression and tunneling progression. The benign tumor fraction *p* determines the progression pattern. We do not model the transition from benign tumors to malignant tumors since we are only interested in the type of progression (dotted arrow).

### Dynamics in the model

In order to describe competition between cells and tumor cell progression, we adopt a Moran model with mutations. This model class has mostly been investigated from a theoretical point of view [18,30,31]. Recently, we applied a Moran model to evaluate tumor regression in pilocytic astrocytoma [19]. Moran models are appropriate to describe a population of fixed size *N* which represents the homeostatic range of competition in our model. The dynamics is as follows. One cell is randomly chosen to undergo cell death and is replaced by the offspring of another chosen cell. During proliferation, a genetic or epigenetic alteration can lead to tumor cell progression. Wild-type cells can progress to benign tumor cells with probability *u* and benign tumor cells progress to malignant tumor cells with probability *v*. We assume that initially all cells are wild-type cells. Hence, the process starts in state 0.

## Analysis of the model

### Choice of spatial cell arrangement

Theoretical studies demonstrated that the interplay between tissue structure, the population size *N* and mutation probabilities *u* and *v* in Moran models are crucial for the dynamics of the model [18,31,32]. In particular, it has been shown that the absorption probability in state *N* on regular structures is the highest if all cells can potentially compete with each other and the lowest for a one-dimensional cell arrangement [18]. Since the tumor-originating cell type is unknown for most cancers also the spatial cell arrangement and realization of competition is unknown [4,33]. Therefore, we consider a space-free and a one-dimensional cell arrangement in order account for this uncertainty by deriving a lower and an upper bound for the absorption probabilities. Figure 2 illustrates the Moran dynamics on these two structures. For the precise definition of the underlying stochastic processes, see Text S1.

**Figure 2:**
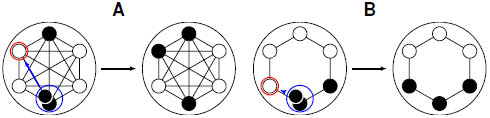
Moran dynamics with different spatial cell arrangements. In the Moran dynamics, a randomly chosen cell proliferates (blue circle) and replaces a neighboring cell which undergoes cell death (red circle). In **A**, the space-free dynamics is illustrated, i.e. each cell can be replaced by any other cell. In **B**, only neighboring cells can be replaced representing a one-dimensional cell arrangement.

### Tumor progression patterns in the model

Three parameter regimes within the model can be distinguished with respect to the tumor progression patterns. Within the *sequential fixation* regime, the benign tumor cell population is primarily able to reach size *N* before a benign tumor cell progresses to a malignant tumor cell. This regime corresponds to primarily sequential progression on the tissue scale. In the *tunneling* regime [30] a malignant clone will occur before the benign population is able to reach size *N* which corresponds to primarily tunneling progression in the model. In the *borderline* regime [32] both sequential fixation and tunneling are possible corresponding to both progression types on the tissue scale. An asymptotic classification of the model behavior with respect to these parameter regimes for large *N* has been theoretically derived in a space-free model [34] and in lattice-like cell arrangements [31]. We showed in a previous work that the exact progression pattern described by the absorption probability in state *N* in the space-free model solely depends on the so-called risk coefficient 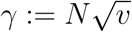 [19]. Analogously, we derive here that the absorption probability in the one-dimensional model solely depends on a one-dimensional risk coefficient 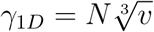.

### Choice of the parameter regime

In order to describe the frequency of sequential tumor formation, we derive the absorption probability of the underlying stochastic process in state *N*. We assume a parameter regime in which no additional mutations from wild-type cells to benign tumor cells occur as long as benign tumor cells are present in the system. The biological consequence of this assumption is that tumors form from the progeny of a single cell [35]. Formally, in the space-free model

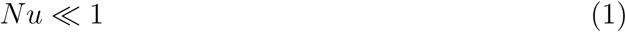

and in the 1*D* model

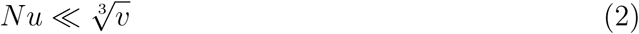

ensures this model behavior, see [18]. For *u* = *v* = 10^−6^, this assumption is valid if *N* ≪ 10^6^ in the space-free model and *N* ≪ 10^4^ in the 1D model.

### Absorption probabilities

In [19], we have shown that the absorption probability in state *N* in the space-free model is given by

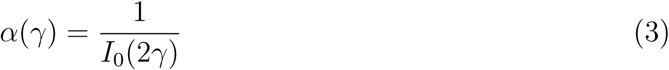

where 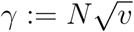 and *I*_*n*_, *nϵ* ℕ_0_, denote the modified Bessel functions of the first kind, see [36]. We refer to the parameter *γ* as *space-free risk coefficient* because it determines the fraction of sequential progression

In this work, we derive the absorption probability in state *N* of the 1*D* model. For this purpose, we utilize first step analysis in order to obtain a linear system of equations for the absorption probabilities in state *N* starting the process with *k,* 1*≤ k ≤N*, benign tumor cells. Subsequently, Cramer’s rule allows to derive the absorption probability starting with one benign tumor cell. Here, it is necessary to calculate two determinants and one of them is approximated by solving a second order difference equation with non-constant coefficients. Finally, we can approximate the absorption probability in state *N* of the 1*D* process starting with a single benign tumor cell as

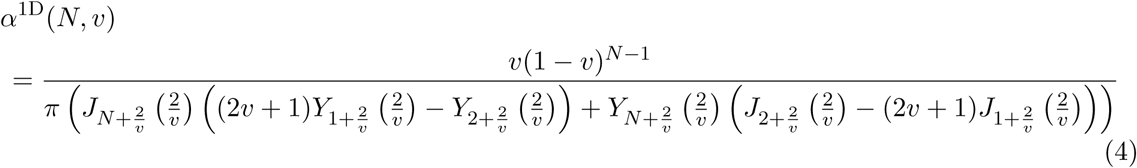

where *J* denotes a Bessel function of the first kind and *Y* a Bessel function of the second kind, see [36]. A detailed derivation of equation (4) is provided in Text S1.

Numerical analysis of the 1D model indicates that the absorption probability in state *N* solely depends on the *1D risk coefficient* given by 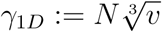, see Figure S5. A detailed derivation of equation (4) and simulation results of the 1D model showing good accordance to this approximation is provided in Table S2.

## Results

### The range of competition determines tumor progression patterns

Our analysis allows to determine the progression patterns in both the space-free and the one-dimensional model in dependency of the competition range *N*. Interestingly, we find that a considerably small value of *N* corresponds to primarily tunneling progression in both the space-free and one-dimensional model. Moreover, the estimates of the parameter *N* largely depend on the considered underlying spatial cell arrangement. In particular, the smaller the number of neighboring cells, the smaller is the estimated competition range. The estimated values for a mutation probability *v* = 10^−6^ per cell division [37] are summarized in Table 1 and visualized in Figure 3. Note that these conclusions also hold for other values of *v* although a smaller value of *v* would increase and a larger value of *v* would decrease the estimates, see Table S3.

**Table 1:** Tumor progression patterns in dependency of the spatial cell arrangement in the model. For these estimates, we have chosen *v* = 10^−6^. Primarily sequential and primarily tunneling progression patterns refer to a fraction of 99.9% of sequential and tunneling progression, respectively.

| progression patterns | 1D model | space-free model |
| --- | --- | --- |
| primarily sequential | $N \leq 17$ | $N \leq 29$ |
| both sequential and tunneling | $17 < N \leq 528$ | $29 < N \leq 4530$ |
| primarily tunneling | $N > 528$ | $N > 4530$ |

**Figure 3:**
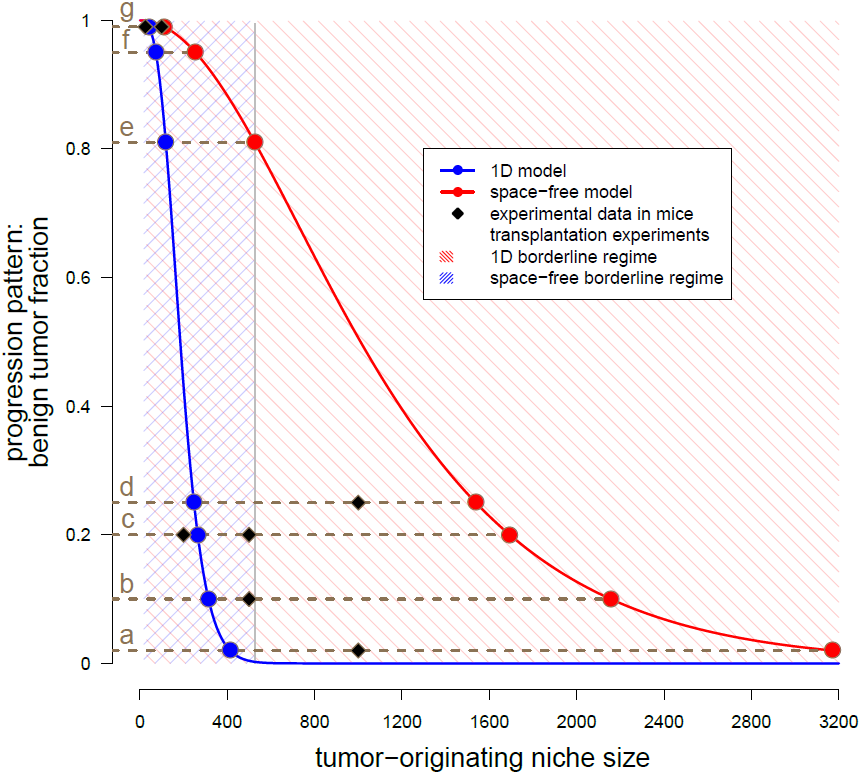
Estimated tumor-originating niche sizes based on tumor progression patterns. This plot shows the benign tumor fraction in the space-free (red) and one-dimensional (blue) model as function of the tumor-originating niche size. The blue curve has been numerically evaluated, see equation [4]. The red curve represents the plot of equation (3). The shaded areas illustrate the borderline parameter regimes, i.e. in which both sequential and tunneling progression are possible for the space-free and the 1*D* model, see Table 1. The dots indicate the estimated tumor-originating niche size and the squares represent experimental data for different tissues, see Table 2 for the values.

### Fate of tumor development is decided in small tissue-specific tumor-originating niches

Our model allows to estimate the range of cellular competition *N* in different human tissues. For these estimations, we calibrate the space-free and 1*D* model with epidemiological data on the diagnosed fraction of benign and malignant tumor subtypes. We performed an extensive literature research to obtain these data which allow to estimate the risk coefficients 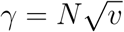 and 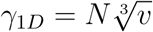 in the following way. We equal the clinically diagnosed fraction of benign tumors *p* with the absorption probabilities in state *N* given by equations [3] and [4] in the Material and Methods section. Subsequently, we numerically calculate the risk coefficients by evaluating the inverse of the absorption probability function at the diagnosed fraction of benign tumors *p*, i.e. *γ* = *α*^−1^(*p*). The resulting estimates of the competition ranges in various tissues are provided in Table 2 and visualized in Figure 3. Our model predicts that the range of competition is considerably small compared to the overall number of cells in a tumor. Note that we do not assume any upper bound for the parameter *N* in our model. Moreover, although the estimates are considerably small, the range of competition largely depends on the tissue. For example, the estimated competition range within the liver is 383 – 2837 cells whereas the estimates for the bone marrow are 18 – 31 cells. Importantly, the estimate of the tumor-originating niche size for the human colon assorts well with the stem cell niche size in colonic crypts of about 40 cells [38] but surely less than 100 cells [39]. Overall, these results can be interpreted as existence of a tissue-specific tumor-originating niche in which the fate of tumor development is decided long before a tumor becomes detectable. Based on our results, we propose that a tumor-originating niche size of 291 – 1928 cells within the human brain might be responsible for the clinically observed fraction of primary and secondary glioblastoma, see Table 2b).

**Table 2:**
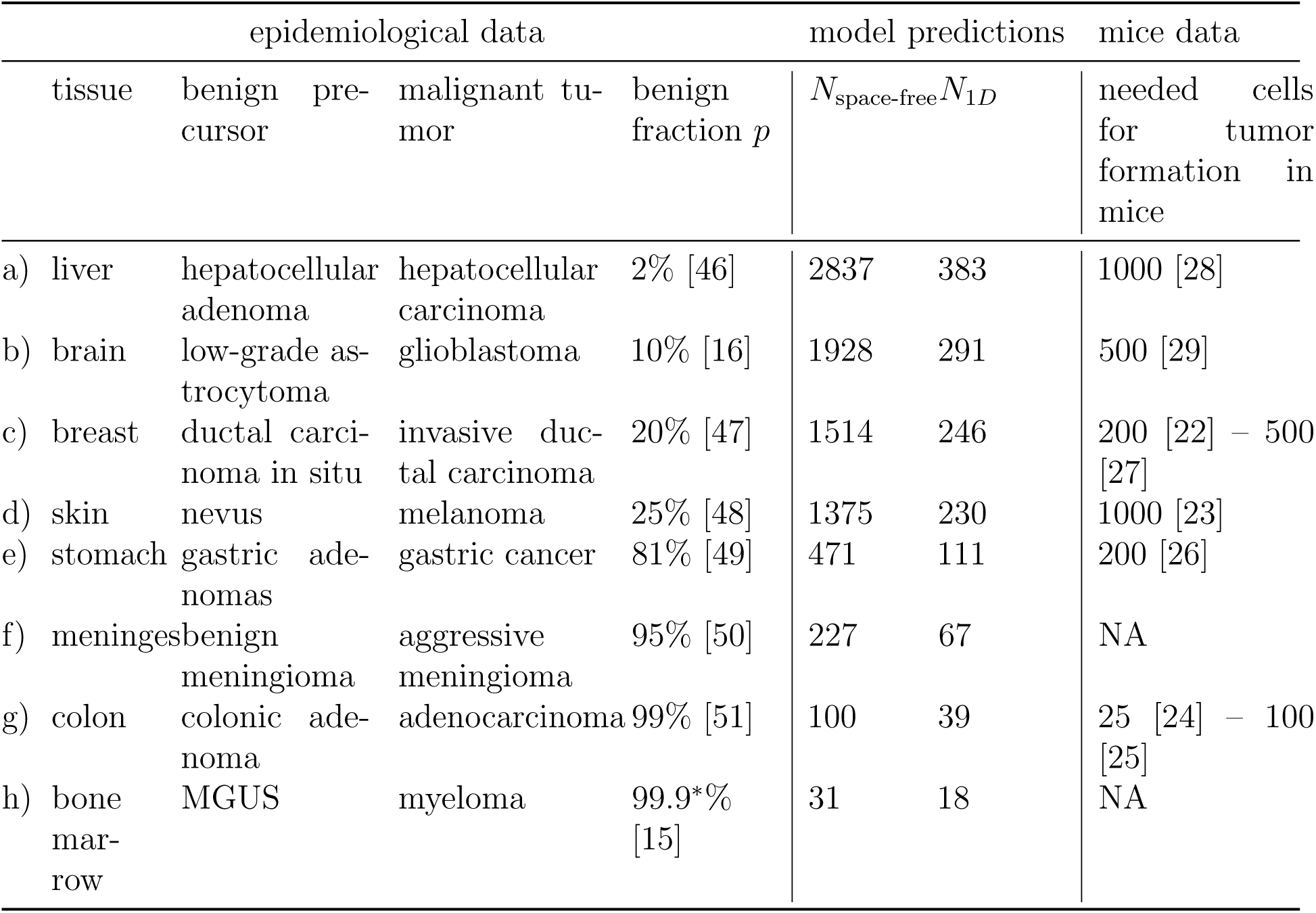
Estimation of the homeostatic competition range *N* in different tissues. This table summarizes epidemiological data on benign and malignant tumor subtypes. We calibrate the model such that the absorption probabilities in state *N* given by equations (3) and (4) are equal to the diagnosed benign tumor fraction *p*. We assume a mutation probability of *v* = 10^−6^ throughout these calculations. The last column contains the necessary number of injected cells to obtain a tumor in mice transplantation experiments. The estimates and mice data are also illustrated in Figure 3. ^*\**^ = estimated, MGUS=monoclonal gammopathy of undetermined significance.

### Predicted tumor-originating niche size agrees well with mice injection experiments

Interestingly, the estimated tumor-originating niche sizes in Table 2 correspond very well to the necessary cell numbers for tumor induction in mice experiments [22–29], see also Figure 3. This observation supports two of our model assumptions. First, the possibility of tumor cell extinction for too small populations in these experiments shows that tumor and wild-type cells compete with each other. Second, the experimental data justify our model assumption that a sufficient number of tumor cells is needed to induce tumor formation on the tissue scale.

## Discussion

On the tissue scale, one observes tumor progression types with and without detectable benign precursor stages. Data on the progression patterns with respect to the ratios of these progression types exhibit large differences between tissues. The underlying cellular processes causing these progression patterns are hardly observable and remain unclear. In this work, we shed light on the early phase of tumor development on the cellular scale with the help of a stochastic model. Our model is based on competition between wild-type and tumor cells and assumes that a sufficient amount of tumor cells is needed for tumor formation. We estimate this number by fitting the model to data on the diagnosed ratios of benign and malignant tumor subtypes. Our model predicts that this number is considerably small compared to the overall number of cells in a clinically detectable tumor and largely depends on the tissue which can be interpreted as existence of a tissue-specific tumor-originating niche. Hence, our results suggest that the fate of tumor development is decided long before a tumor becomes detectable.

Interestingly, our estimates of the tumor-originating niche size of about 39 cells for colon cancer agrees well with the number of stem cells found in one colonic crypt [38]. Indeed, it is the current understanding that colon adenomas and carcinomas develop within one colonic crypt with intestine stem cells likely to be the cell type of origin [40]. This demonstrates that our model might be utilized to predict tumor-originating niche sizes, thereby allowing to infer the potential cell type of tumor-origin for cancers from other tissues in which the origin is still elusive, e.g. for glioblastoma [41]. Glioblastoma can be divided into two classes dependent on the progression dynamics. In about 90% of cases, glioblastoma occur *de novo*, i.e. without evidence of a less malignant precursor lesion (primary glioblastoma) whereas 10% develop slowly by progression from low-grade gliomas (secondary glioblastoma). Using this data, our model predicts that the size of the tumor-originating niche from which glioblastoma develop is about 291 – 1928 cells. Neural stem cells (NSCs) of the subependymal zone (SEZ) have been suggested as a potential cell of origin for glioblastoma. Moreover, recent experimental evidence regarding NSCs in the SEZ of the adult brain suggests that the total number and fate of NSCs is regulated by a density-dependent mechanism [42]. Importantly, the finding in [42] that the fate of a NSC, e.g. activation or quiescence, is coupled to its neighbors perfectly fits to our hypothesis of cells competing within a certain range. Interestingly, the authors also suggest that the fate of active NSCs is coupled to the total number of neighboring NSCs in a shared locally restricted area which suggests that this area is a potential candidate for the tumor-initiating niche in the adult brain. It would be interesting to investigate if the range of coupled NSCs fits to our predicted size of the tumor-originating niche for glioblastoma.

From a modeling perspective, our analysis allows to distinguish different model regimes in Moran models not only asymptotically as in [18], but also for finite values of the system size *N*, see Table S3. and Figure S5. We find that the risk coefficient in both the space-free and the 1*D* model does not only distinguish the different model regimes, but quantifies the proportion of tunneling and sequential progression. Moreover, we show that the corresponding asymptotic results in dependency of the risk coefficient provide very good approximations even for small population sizes *N*. This finding obviates the need of ad-hoc rules like *a≪ b* if *a/b≤*1*/*10 to choose the appropriate parameter regime [43].

The potential existence of tumor-originating niches in which tumor fate is decided at an early stage of the cellular multistep process supports the view that cancer development is an ecological process [44,45]. Ecology studies the dynamics of communities of species and their interactions and describe the origin of new species. From this point of view, the size of the tumor-originating niche might represent a critical effective population size that has to be reached by the progeny of the tumor-originating cell type in order to establish a tumor on the tissue scale. A deeper understanding of the processes and the origin of the tumor-originating niche contributes to the understanding of the early phase of tumor development. Further modeling and experimental effort is needed to understand this early phase of tumor development on the cellular scale in a better way. In this work, we demonstrated how observable quantities on the tissue scale might be utilized to achieve this goal.

## Supplementary Material

**S1 Text**

The supplementary text contains the precise definitions of the Moran models and detailed analytical derivations.

**S2 Table**

Comparison of the analytical approximation given and simulation results from 10000 trajectories of the underlying stochastic process of the one-dimensional model for the probability of absorption in state N. The results are also visualized in Figure S5

**S3 Table**

This table summarizes the risk coefficient regimes with respect to the different progression patterns of the model. Primarily sequential and primarily tunneling progression patterns refer to a fraction of 99.9% of sequential and tunneling progression, respectively.

**S4 Table**

This table summarizes regimes for the parameter *N* with respect to the different progression patterns of the models in dependency of the mutation probability *v*. Primarily sequential and primarily tunneling progression patterns refer to a fraction of 99.9% of sequential and tunneling progression, respectively.

**S5 Figure**

We numerically approximated the absorption probability in state *N* for different values of *N* and *v* such that the risk coefficient *γ*_1D_ is constant. This analysis suggests that the absorption probability solely depends on the risk coefficient *γ*_1D_ for approximately *N*≥40. The squares indicate the results of simulation studies of the absorption probability in state *N* and therefore the benign tumor fraction in the model, see also Table S2

## Acknowledgments

The authors thank the Center for Information Services and High Performance Computing at TU Dresden for providing an excellent infrastructure.

